# Loss of mutually protective effects between osteoclasts and chondrocytes in damaged joints drives osteoclast-mediated cartilage degradation via matrix metalloproteinases

**DOI:** 10.1101/2021.01.03.425116

**Authors:** Quitterie C Larrouture, Adam P Cribbs, Sarah J Snelling, Helen J Knowles

**Affiliations:** Department of Pathology, University of Pittsburgh, Pittsburgh, USA; Botnar Research Centre, Nuffield Department of Orthopaedics, Rheumatology and Musculoskeletal Sciences, University of Oxford, OX3 7LD, UK

## Abstract

Osteoclasts are large multinucleated cells that resorb bone to regulate bone remodelling during skeletal maintenance and development. It is overlooked that osteoclasts also digest cartilage during this process, as well as in degradative conditions including osteoarthritis, rheumatoid arthritis and primary bone sarcomas such as giant cell tumour of bone. This study explores the poorly understood mechanisms behind the interaction between osteoclasts and cartilage. Morphologically, osteoclasts differentiated on acellular human cartilage formed multinucleated cells expressing characteristic osteoclast marker genes (e.g. *CTSK, MMP9*) and proteins (TRAP, VNR) that visibly damaged the cartilage surface by SEM, but without the formation of resorption pits. Osteoclasts caused increased glycosaminoglycan (GAG) release from acellular and cellular human cartilage that was dependent on direct contact with the substrate. Direct co-culture with chondrocytes during osteoclast differentiation increased the number of large osteoclasts formed. When osteoclasts were cultured on dentine, direct co-culture with chondrocytes inhibited osteoclast formation and reduced basal degradation of cartilage. This suggests a mutually protective effect on their ‘native’ tissue between bone-resident osteoclasts and chondrocytes, that is reversed when the joint structure breaks down and osteoclasts are in contact with non-native substrates. Mechanistically, osteoclast-mediated cartilage degradation was inhibited by the pan-MMP inhibitor GM6001 and by TIMP1, indicative of a role for soluble MMPs. RNA sequencing and RT-qPCR analysis identified *MMP8* as overexpressed in osteoclasts differentiated on cartilage versus dentine, while *MMP9* was the most highly expressed MMP on both substrates. Inhibition of either *MMP8* or *MMP9* by siRNA in mature osteoclasts reduced GAG release, confirming their involvement in cartilage degradation. Immunohistochemical expression of MMP8 and MMP9 was evident in osteoclasts in osteosarcoma tissue sections. Understanding and controlling the activity of osteoclasts might represent a new therapeutic approach for pathologies characterized by cartilage degeneration and presents an attractive target for further research.

## Introduction

Osteoclasts are large multinucleated cells that resorb bone. They are formed by the fusion of CD14+ monocytes and regulate skeletal development and maintenance via homeostatic interactions with bone-forming osteoblasts.

It is often overlooked that osteoclasts also digest cartilage. During skeletal development, endochondral ossification replaces cartilage templates of long bone with mineralized bone following osteoclast-mediated digestion of the calcified cartilage matrix secreted by hypertrophic chondrocytes [1-3]. Osteoclasts also degrade the calcified cartilaginous callus during bone fracture healing. In a murine model, mice deficient in osteoprotegerin (OPG), a negative regulator of osteoclasts, degrade the cartilaginous callus faster, with increased osteoclast numbers and faster union of the fractured bone [4].

Dysregulation of the balance between osteoclasts and osteoblasts towards relative osteoclast overactivation results in pathological osteolysis in conditions from osteoporosis and bone cancer to rheumatoid arthritis (RA). RA, a connective tissue disease of the joints, is also characterised by cartilage destruction. Inflammatory cytokines activate fibroblast-like synoviocytes (FLS) and T cells to produce osteoclastogenic macrophage colony stimulating factor (M-CSF) and receptor activator of nuclear factor kappa B ligand (RANKL), promoting the fusion of monocytes to form multinucleated osteoclasts. These osteoclasts drive joint destruction via physical contact with bone and both non-mineralised and calcified cartilage [5,6]. Osteoclasts also invade articular cartilage in human knee osteoarthritis (OA) [7]; OA being the most common disease of cartilage loss.

Inflammatory cytokines such as TNFα potentiate the osteoclastogenic effect of RANKL [8], which could explain how TNFα-blocking agents protect against bone and cartilage loss in patients with RA [9]. In human TNF-transgenic mice, which develop destructive arthritis associated with enhanced formation of osteoclasts, both an anti-TNFα antibody and the osteoclast-inhibiting bisphosphonate zoledronate block bone erosion and cartilage damage and reduce osteoclast formation within the inflamed synovium [10]. Denosumab, a monoclonal anti-RANKL antibody, inhibits osteoclast formation, suppresses bone resorption and causes a reduction in the cartilage turnover marker urine CTX-II/creatinine in patients with RA [11]. Osteoclast-inhibiting bisphosphonates [12,13] and OPG [14] are also chondroprotective in murine models of OA.

Mature osteoclasts have a morphology highly specialised for bone-resorption [15]. Resorption is preceded by osteoclast attachment to the bone matrix, mediated by integrins such as the vitronectin receptor (VNR, αvβ3 integrin). This is followed by polarization and cytoskeletal reorganization; F-actin-rich podosomes form a dynamic actin ring which, alongside the integrins, forms a sealing zone that isolates the highly folded bone-facing ruffled border membrane from the extracellular environment [15]. Ultrastructural features of osteoclasts resorbing calcified cartilage exhibit many features common to bone-resident osteoclasts including abundant mitochondria, vacuolation, lysosomes and deep infoldings at points of contact with the calcified matrix [3].

Osteoclasts on bone and those on cartilage (sometimes termed ‘chondroclasts’ [16]) also have similar basic molecular characteristics. Multinucleated cells form within the inflammatory synovium in RA but only express the calcitonin receptor, a marker of fully differentiated osteoclasts, when they come into direct contact with calcified cartilage or subchondral bone, suggesting that contact with either form of mineralised tissue can direct the final stages of differentiation [17]. Multinucleated cells digesting cartilage in RA, OA and the primary bone tumour giant cell tumour of bone (GCTB) express an osteoclast phenotype (CD14−, HLA-DR−, CD45+, CD68+ CD51+, TRAP+, cathepsin K+ and MMP9+) [18].

However, the mechanism of osteoclast formation on and degradation of cartilage is poorly understood. Osteoblasts produce the necessary cytokines for homing osteoclastogenesis to bone, but chondrocytes can also produce these factors [19,20]. CD14+ monocytes differentiated on healthy human articular cartilage express immunophenotypic markers characteristic of osteoclasts (CD68+, CD14− and CD51+) and degrade cartilage via release of glycosaminoglycans (GAG). Primary human osteoclasts from GCTB and pigmented villonodular synovitis (PVNS) tissue can also release GAG from cartilage [18]. Maintenance of articular cartilage is a balance between anabolic and catabolic pathways with degradation mediated in large part by the matrix metalloproteinases (MMPs), which can also be produced by osteoclasts [7,21].

Despite evidence of a physical interaction between osteoclasts and cartilage in RA, OA and GCTB the mechanism(s) by which osteoclasts degrade cartilage remain unknown. The aim of this study was to enhance our understanding of the interaction between osteoclasts and cartilage; to ascertain whether osteoclasts degrade cartilage using the same cellular machinery as for bone resorption and whether chondrocytes affect this process. If osteoclasts contribute to driving cartilage degradation, understanding and controlling their activity would represent an important perspective for diseases characterized by degeneration of cartilage.

## Materials and Methods

### Ethics

Use of leucocyte cones for osteoclast differentiation was approved by the London-Fulham Research Ethics Committee (11/H0711/7). Archival pathological bone and joint specimens were obtained from the Nuffield Orthopaedic Centre, Oxford, UK. Cartilage was obtained at the time of surgery from patients undergoing total knee arthroplasty for OA at the Nuffield Orthopaedic Centre. Samples were obtained via the Oxford Musculoskeletal Biobank and were collected with informed donor consent in full compliance with national and institutional ethical requirements, the United Kingdom Human Tissue Act and the Declaration of Helsinki (HTA Licence 12217, Oxford REC C 09/H0606/11+5, London-Fulham Research Ethics Committee 07/H0706/81).

### Osteoclast differentiation

Peripheral blood mononuclear cells were isolated from human leucocyte cones (NHS Blood and Transplant, Bristol, UK) by density gradient centrifugation. CD14+ monocytes were positively selected using MACS CD14+ microbeads (Miltenyi Biotech, Surrey, UK) and seeded at 0.25 × 10^6^ cells/well into 96-well plates containing dentine discs, at 1 × 10^6^ cells/well into 48-well plates containing cartilage pieces, or at 1 × 10^6^ cells/well into 24-well plates. After overnight adhesion, dentine discs and cartilage pieces were transferred to new wells. Osteoclast differentiation was induced by treatment with 25 ng/ml M-CSF (R&D Systems, Abingdon, UK) and 50 ng/ml RANKL (Peprotech, London, UK) in α-MEM containing 10% FBS, 50 IU/ml penicillin, 50 μg/ml streptomycin sulphate and 2 mM L-glutamine. Differentiation medium was refreshed every 2-3 days for 10-13 days.

For inhibitor experiments, mature osteoclasts were treated with bafilomycin (Cell Signalling Technology, Hitchin, UK), E64 (Cambridge Bioscience, Cambridge, UK), GM6001 (Selleckchem, Munich, Germany) or recombinant human TIMP1 (R&D Systems).

### Preparation of dentine and cartilage substrate

Dentine (elephant ivory discs; HM Revenue & Customs, Heathrow Airport, UK) was prepared by cutting 250 μm transverse wafers using a low-speed saw and diamond-edged blade (Buehler, Coventry, UK), out of which 4 mm diameter discs were punched. Cartilage tissue was washed to remove blood and debris and cut to form approximately 6×6 mm squares. To generate acellular cartilage, explants were stored at −80°C then thawed and snap-frozen in liquid nitrogen to ensure killing of the chondrocytes. Cellular cartilage pieces (containing live chondrocytes) were maintained in αMEM for 4 days prior to experimental use. Alamar Blue was used to confirm that chondrocytes in cellular cartilage were metabolically active compared to those in acellular cartilage pieces.

### Histology and immunohistochemistry

Osteoclasts cultured on dentine or cartilage were fixed in 4% formalin. Dentines were decalcified in 0.5M EDTA prior to paraffin embedding. H&E staining was performed on transverse 5 μm sections. For immunohistochemistry, antigen retrieval of deparaffinised OA tissue sections was performed by immersion in hot citric acid solution. Sections were exposed to primary rabbit monoclonal antibodies against MMP-8 (ab81286, 1:1000, Abcam, Cambridge, UK) or MMP-9 (ab73734, 1:1000, Abcam) overnight at 4C or a PBS control and staining was visualised with DAB. Image acquisition was performed using a Zeiss AxioImager MI microscope, AxioCam HRC camera and AxioVision software. Osteoclasts in tissue sections were considered as large, multinucleated cells containing ≥3 nuclei; a widely accepted identification criterion [18,22,23].

### Characterisation of osteoclasts

Tartrate-resistant acid phosphatase (TRAP) staining of osteoclasts was performed on formalin-fixed cells using naphthol AS-BI phosphate as a substrate and reacting the product with fast violet B salt at 37°C for 3 h. TRAP-positive multinucleated cells containing 3≥ nuclei were considered as osteoclasts. Expression of the vitronectin receptor (VNR, CD51/61, αVβ3-integrin), a marker of mature osteoclasts, was determined by immunohistochemistry using a murine monoclonal antibody targeting CD51/61 (MCA757G, 1:200; AbD Serotec, Oxford, UK) and visualised with DAB. Immunofluorescent staining of F-actin filaments was performed on formalin-fixed cells for 30 min using TRITC-conjugated phalloidin (P5282, 1:200; Sigma-Aldrich, Poole, UK), mounted with fluoroshield containing DAPI (Sigma)and visualised at 450-480nm. Image acquisition was performed using a Zeiss AxioImager MI microscope, AxioCam HRC camera and AxioVision software. Quantification was performed using ImageJ software (National Institutes of Health, Bethesda, USA) to count the average number of osteoclasts per field of view.

### RT-PCR

RNA was extracted in TRIzol (Invitrogen), purified (Direct-Zol RNA miniprep kit; Zymo Research, Cambridge, UK) and reverse transcribed (High capacity cDNA reverse transcription kit; Applied Biosystems, California, USA). Quantitative PCR was performed using Fast SYBR Green Master Mix or TaqMan Fast Advanced Master Mix in a Viia7 Real-Time PCR system (Applied Biosystems, Warrington, UK). Human primers were either pre-validated Quantitect primers against *MMP1* (Hs_MMP1_1_SG), *MMP3* (Hs_MMP3_1_SG), MMP9 (Hs_MMP9_1_SG), *MMP13* (Hs_MMP13_1_SG) or *ACTB* (Hs_ACTB_2_SG) (Qiagen, Manchester, UK), Primerdesign primers against *MMP8* (JN215643) or *ADAMTS9* (JN171532) (Primerdesign, Chandlers Ford, UK) or designed in-house (Supplementary Table 1). A gene was not considered to be expressed when the average cycle threshold (Ct) value per condition was higher than 34. The expression of individual mRNAs was calculated relative to expression of β-actin (*ACTB*) using the (2-ΔCt) method.

### Scanning Electron Microscopy

Dentine slices and cartilage pieces were fixed in 4% formalin, sputter-coated with gold using the SC7620 Mini Sputter Coater System (Quorum Technologies, Lewes, UK) and imaged using a Philips XL 30/SEM with field emission gun.

### Quantification of GAG and collagen release

The 1,9-dimethylmethylene blue (DMMB) assay quantifies sulphated GAGs (chondroitin 4- and 6-sulphate, heparin, keratin and dermatan sulphates). Conditioned media was diluted in NaH_2_PO_4_ / Na_2_HPO_4_ buffer solution (pH 6.5). Diluted sample (40 μl) was mixed with 250 μl DMMB solution (16 mg DMMB, 50 mM glycine, 40 mM NaCl, 0.1 M HCl, pH 3) and the absorbance was measured at 530 nm against a chondroitin sulphate standard curve.

Collagen was quantified by measuring hydroxyproline using the dimethylaminobenzaldehyde (DAB) and chloramine T assay. Conditioned media was hydrolysed with an equal volume of concentrated HCl for 18 hr at 105°C. The precipitate dried, resuspended in dH_2_O, and 40 μl was reacted with 25 μl chloramine T solution (140 mg chloramine T, 2 ml dH_2_O, 8 ml acetate-citrate buffer) for 4 min to oxidise free hydroxyproline. This was incubated with 150 μl DAB solution (30 ml propan-2-ol, 10 ml DAB solution [20g DAB, 30 ml 70% perchloric acid]) at 65°C for 35 min. Absorbance was measured at 560 nm and compared with a hydroxyproline standard curve.

### Isolation of chondrocytes

Chondrocytes were isolated from cartilage tissue by overnight digestion with collagenase (1 mg/ml) then passed through a 70 μm cell strainer (Falcon Becton Dickinson, Oxford, UK). Chondrocytes were grown to 80% confluence and used for experiments up to passage 3.

### Co-culture experiments

Direct co-culture was performed with CD14+ monocytes (1 × 10^6^ / well) and chondrocytes (4 × 10^4^ / well) seeded in 24-well plates in α-MEM. For Transwell experiments, chondrocytes (4 × 10^4^) were seeded onto 0.4 μm filter cell culture inserts. After 24 h the Transwells were moved into 24-well plates in which CD14+ monocytes (1 × 10^6^ / well) had been seeded on cell culture plastic or dentine discs in the lower chamber. For Transwell cartilage experiments, cartilage was placed in the cell culture insert, with monocytes in the lower chamber. Cultures were treated for 9-10 days with M-CSF and RANKL as for osteoclast differentiation.

### RNAseq

RNA was DNase I treated, cleaned and concentrated (Zymo RNA Clean and Concentrator, Zymo Research) then enriched for poly(A) mRNA (NEBNext poly(A) mRNA Magnetic Isolation Module, NEB Biosystems, Ipswich, UK). Sequencing libraries were prepared using the NEBNext Ultra II RNA Library Prep kit (New England Biolabs). RNA and DNA quality were assessed using High Sensitivity RNA or DNA Screentape and an Agilent 4200 tapestation. Single-indexed and multiplexed samples were run on an Illumina Next Seq 500 sequencer using a NextSeq 500 v2 kit (FC-404-2005; Illumina, Can Diago, CA) for paired-end sequencing.

A computational pipeline was written calling scripts from the CGAT toolkit https://github.com/cgat-developers/cgat-flow) [24,25]. Sequencing reads were de-multiplexed based on the sample index and aligned to the human genome assembly version 38 (GRCh38) using the STAR (Spliced Transcripts Alignment to a Reference) aligner [26] At least 14 million aligned reads were obtained per sample. Reads were mapped to genes using featureCounts v1.4.6 (part of the subreads package), in which only uniquely mapped reads were counted to genes. Differential expression analysis was performed using DESeq2 [27] between three groups: osteoclasts cultured on cell culture plastic, dentine or cellular cartilage.

### Gelatin zymography

Zymography was used to assess the activity of different MMP isoenzymes. Proteins are separated in sodium dodecyl sulphate (SDS). After 9 days of osteoclast differentiation on acellular cartilage, the media was changed to FBS-free media with M-CSF and RANKL for 24 h. Conditioned media was collected, centrifuged to remove cell debris, and mixed with 2 volumes of Zymogram sample buffer (1610764; Bio-Rad). Samples were loaded onto a 10% acrylamide gel containing copolymerised gelatin (ZY00100BOX; ThermoFisher Scientific) run at 150 V for approximately 2 h. Active recombinant human MMP8 (908-MP-010, R&D systems) and MMP9 (911-MP-010, R&D systems) were used as positive controls. Gels were washed in renaturation buffer (1610765, BioRad) to remove SDS and then in development buffer (50 mM Tris, 10 mM CaCl2, 50 mM NaCl, 0.05 % Brij 35, pH 7.6) to remove renaturation buffer and activate MMPs. Gels were then processed for silver staining and scanned using an Epson V700 Photo scanner and Epsonscan software.

### siRNA targeting *MMP8* and *MMP9*

Osteoclasts were differentiated on cell culture plastic for 7 days before transfection with two separate siRNAs targeting either *MMP8* (105332, 112911; Life Technologies) or *MMP9* (104072, 113182; Life Technologies) or a non-specific siRNA control at 25 nM concentration using lipofectamine RNAiMax (Life Technologies). Duplexes were removed after 6 h and replaced with osteoclast differentiation medium.

### Statistics

Statistical analysis was performed in GraphPad Prism 8 (GraphPad Software, California, USA). Statistical significance was determined using one-way ANOVA with Dunnett’s Multiple Comparison or Student’s T-test as appropriate. Results were considered significant at p<0.05. All figures include at least three independent experiments and are plotted as mean ± SD.

## Results

### Osteoclasts can differentiate on cartilage and visibly degrade the cartilage matrix

Multinucleated cells morphologically characteristic of osteoclasts were observed adjacent to visibly eroded mineralised and non-mineralised cartilage in cases of OA, RA and GCTB (Figure 1a). Human osteoclasts differentiated on dentine produced clear resorption pits beneath the cells, visible by both light microscopy and SEM, associated with formation of an F-actin ring (Figure 1b). In contrast, and despite formation of large multinucleated cells expressing the osteoclast markers TRAP and VNR, human osteoclasts differentiated on unmineralised acellular articular cartilage did not form an F-actin ring and did not perform visible erosion by light microscopy (Figure 1c). No difference in the expression of classical osteoclast marker genes (*ACP5* (*TRAP), ATP6V1A 21S, CA2, CLCN7, CSTK, MMP9*) was evident between osteoclasts formed on dentine or on cartilage (Figure 1d). SEM images confirmed that osteoclasts on cartilage were of similar size to those on dentine but were more spherical in appearance. The cartilage surface was visibly damaged around osteoclasts despite no visible resorption tracks (Figure 1e) indicating that osteoclasts do degrade cartilage but, despite having a similar molecular profile, not in the same way as bone matrix.

**Figure 1:**
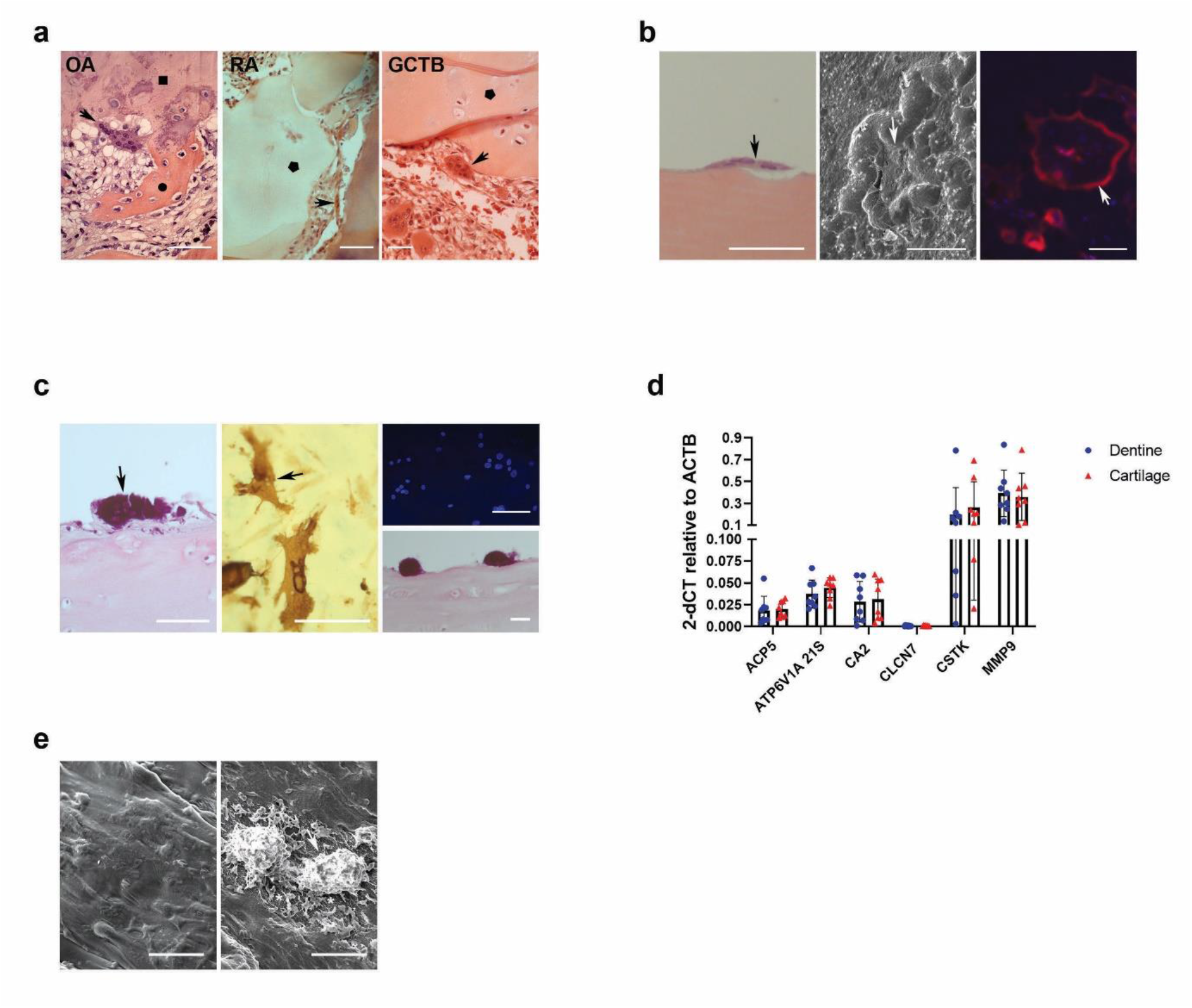
Osteoclasts visibly degrade cartilage matrix. (a) H&E staining of OA knee, RA and GCTB tissue. Multi-nucleated osteoclasts (arrow) resorb subchondral bone 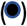 and invade mineralised 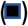 and unmineralised cartilage 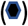. (b) Osteoclasts (arrow) differentiated on dentine with (left) H&E-stained transverse section showing visible subcellular resorption, (middle) SEM image of a 97.49 μm diameter osteoclast surrounded by resorption tracks, and (right) F-actin rings (red) around multi-nucleated (blue) osteoclast. (c) Osteoclasts (arrow) differentiated on acellular cartilage showing (left, top right) TRAP-stained transverse section through multi-nucleated osteoclasts with no visible subcellular resorption and (middle) CD51/61-positive osteoclasts with (bottom right) no evidence of F-actin ring formation. (d) Expression of classical osteoclast marker genes by osteoclasts differentiated on dentine or acellular cartilage. ACP5 (TRAP), ATP6V1A 21S, CA2, CLCN7 (H+/Cl-transporter channel 7), CSTK (cathepsin K), MMP9. *, p<0.05; ***, p<0.0001; n= 8. (e) SEM images of (left) acellular cartilage and (right) osteoclasts (arrow) on acellular cartilage showing visible degradation the surrounding cartilage matrix (*). Scale bars = 100μm.

### Regulation of osteoclast differentiation and function by chondrocytes is dependent on the osteoclast substrate

To quantify resorption of the different substrates, we measured osteoclast-mediated release of collagen and GAG. Generation of active osteoclasts was confirmed by release of collagen from dentine during resorption pit formation (Figure 2a). Osteoclasts did not cause release of collagen from cartilage (Supplementary Figure 1a, b) but did release GAG from both acellular cartilage (chondrocytes killed by freeze-thawing, Supplementary Figure 1c) and cellular cartilage containing live chondrocytes (Figure 2a). Increased GAG release was not evident when osteoclasts were cultured on dentine or plastic with cartilage in a Transwell insert, suggesting that direct contact between osteoclasts and cartilage is essential for proteoglycan degradation (Figure 2b, c). Interestingly, basal GAG release from cellular cartilage was inhibited when distant osteoclasts (i.e. those not in direct contact with cartilage) were cultured on dentine but not plastic (Figure 2c), suggesting a protective effect of bone-resident osteoclasts on the cartilage matrix via effects of secreted products on active chondrocytes.

**Figure 2:**
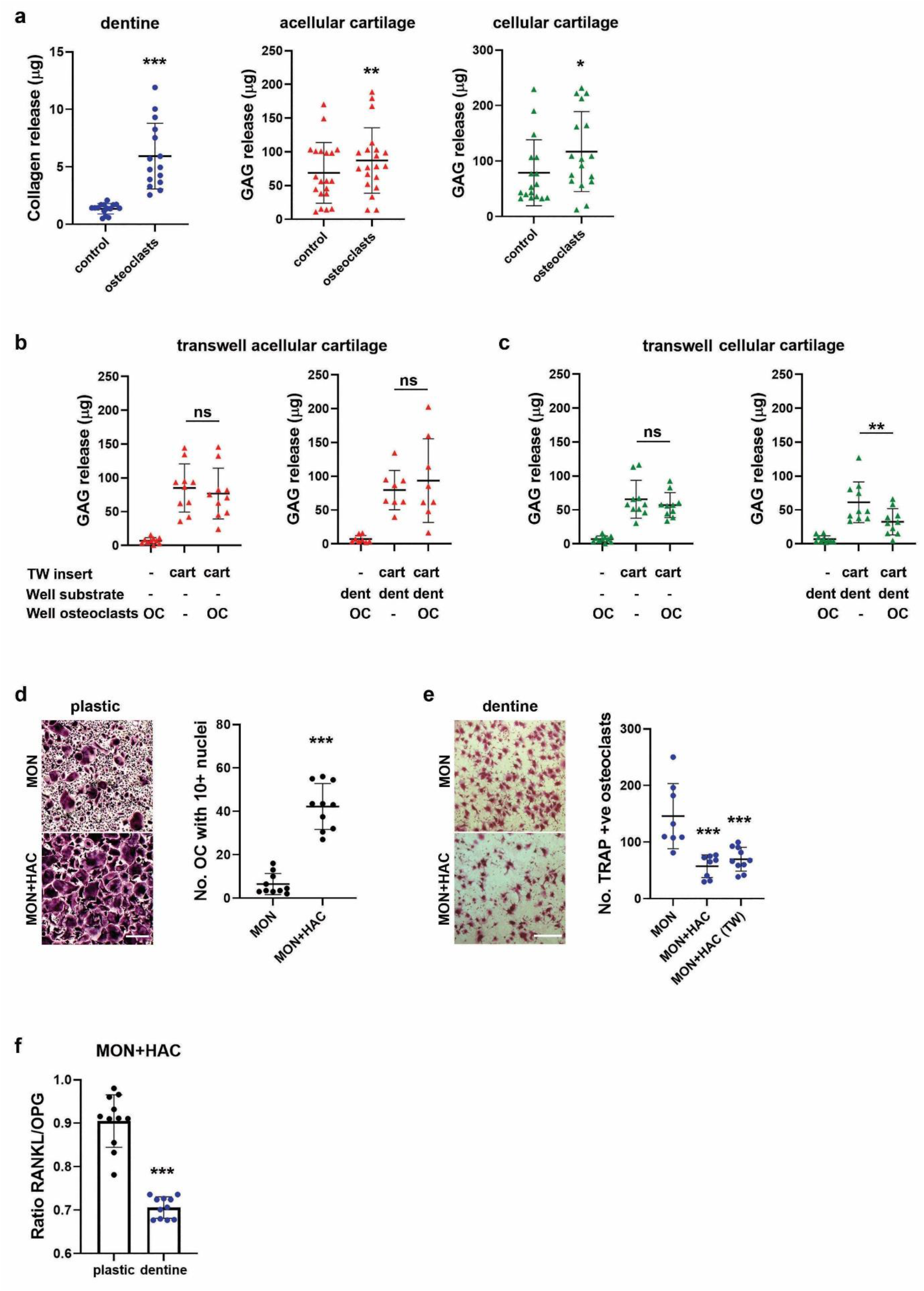
Substrate-dependent regulation of osteoclasts by chondrocytes. (a) Osteoclast-mediated release of (left, n=16) collagen from dentine and GAG from (middle, n=20) acellular or (right, n=17) cellular cartilage after 13 days of differentiation. (b, c) GAG release from indirect co-culture experiments with osteoclasts differentiated on either plastic or dentine for 13 days with pieces of (b) acellular cartilage or (c) cellular cartilage in a Transwell insert. n=10. (d, e) TRAP-staining following 10 days’ differentiation of CD14+ monocytes into osteoclasts by direct co-culture with chondrocytes and exogenous M-CSF and RANKL. Differentiation performed on (d) cell culture plastic (scale bar = 400 μm) with quantification of the number of TRAP-positive cells with ≥10 nuclei, n=10, or (e) dentine discs (scale bar = 500 μm) with quantification of the total number of TRAP-positive cells, n=8. (f) Expression ratio of RANKL:OPG in direct co-cultures of osteoclasts and chondrocytes on plastic or dentine, n=11. *, p<0.05; **, p<0.01; ***, p<0.0001.

Direct co-culture of CD14+ monocytes with chondrocytes increased the number of large osteoclasts (>10 nuclei) formed on plastic (Figure 2d), but reduced the number formed on dentine in a manner independent of direct contact (Figure 2e). This might be partially due to a reduced RANKL:OPG expression ratio in chondrocytes cultured on dentine (Figure 2f). This suggests that factors secreted by chondrocytes inhibit the formation and activity of bone-resident osteoclasts, but that chondrocytes can stimulate osteoclast formation on non-bone sites.

### MMPs drive osteoclast-mediated degradation of cartilage

We next sought to identify the proteinases responsible for osteoclast-driven degradation of cartilage by inhibiting key components of bone resorption. The efficacy of bafilomycin and E64 to inhibit acidification of the resorption lacunae and cathepsin K activity respectively was confirmed in osteoclasts cultured on dentine (Supplementary Figure 2a). Neither inhibitor affected osteoclast-mediated release of GAG from cartilage. However, the pan-MMP inhibitor GM6001 significantly reduced osteoclast-mediated GAG release from both acellular (Figure 3a) and cellular (Figure 3b) cartilage. The greater reduction with cellular cartilage (57% reduction versus 43% in acellular cartilage) was due to inhibition of basal release of GAG from cellular cartilage by GM6001 in the absence of other stimuli (Supplementary Figure 2b), confirming that live chondrocytes release MMPs that degrade cartilage. Recombinant human tissue inhibitor of metalloproteinase 1 (TIMP1) also inhibited osteoclast-mediated release of GAG from acellular cartilage (Figure 3c), establishing the active enzyme as a soluble MMP.

**Figure 3:**
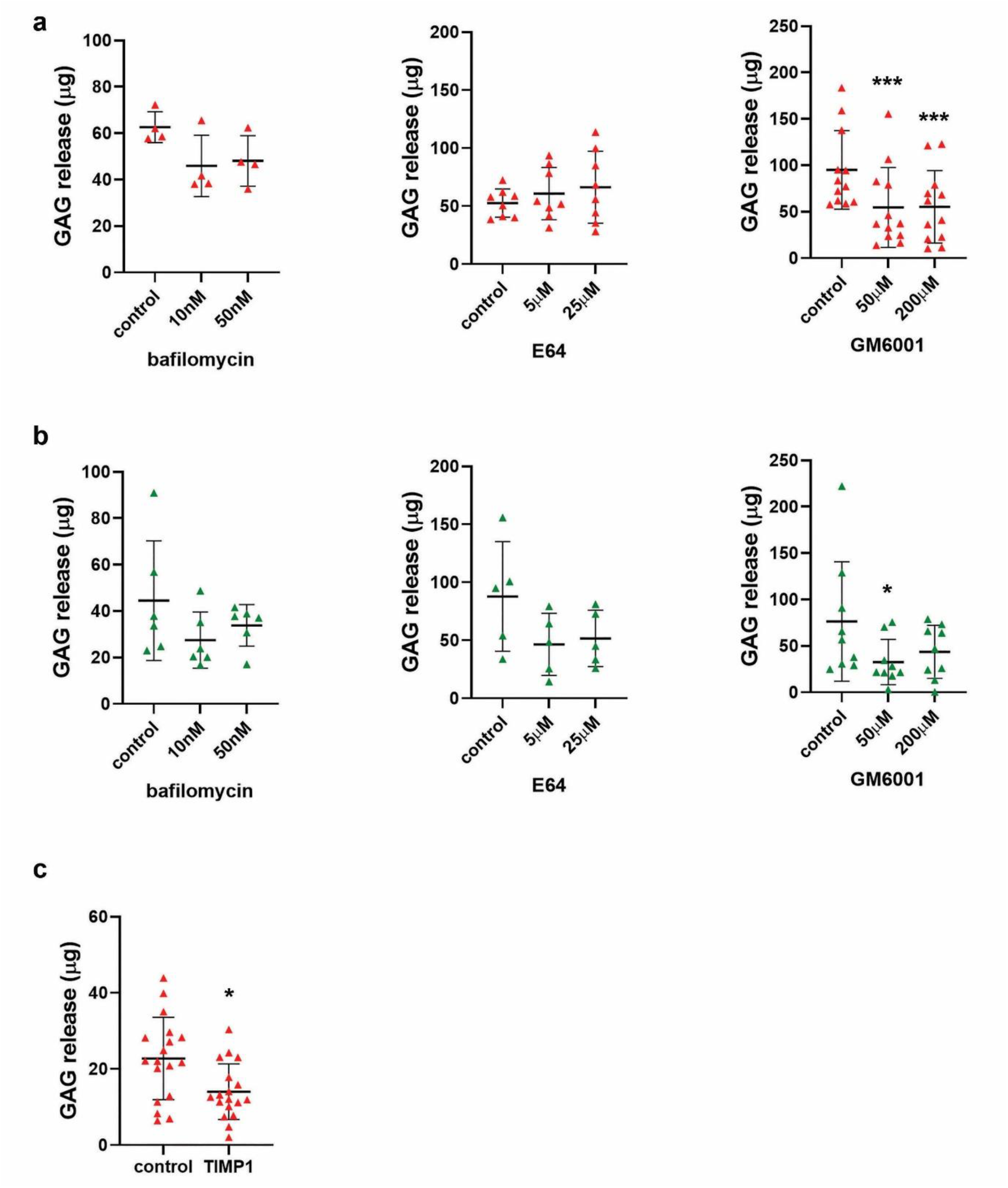
MMPs drive osteoclast-mediated cartilage degradation. (a,b) Effect of bafilomycin, E64 and GM6001 on osteoclast-mediated release of GAG from (a) acellular or (b) cellular cartilage after 13 days of differentiation. N varies for each graph, as represented by the number of data points. (c) Effect of recombinant human TIMP1 (100 nm) on GAG release from osteoclasts cultured on acellular cartilage, n=18. *, p<0.05; ***, p<0.0001.

### MMP8 and MMP9 are the primary MMPs responsible for osteoclast-mediated release of GAG from cartilage

To identify the specific MMP(s) responsible for osteoclast-mediated cartilage degradation, as well as other contributory matrix-degrading genes, we used RNAseq to compare the expression profile of osteoclasts differentiated on cell culture plastic, acellular cartilage and dentine. Osteoclasts on dentine clustered away from the other substrates, suggesting distinct gene expression profiles between bone-resident osteoclasts and those on ‘non-bone’ substrates (Figure 4a). *MMP8* showed the greatest fold upregulation in osteoclasts on cartilage versus dentine (8.89-fold, p = 0.0133) (Figure 4b, Supplementary Figure 3,Table 1). No other of the top regulated genes had functions suggestive of an involvement in cartilage degradation. We therefore explored expression of other members of the large sub-family of soluble MMPs. No difference in expression of any *MMP* was observed between substrates, although *MMP9* mRNA was consistently expressed at a higher level than other *MMPs* (Figure 4c).

**Figure 4:**
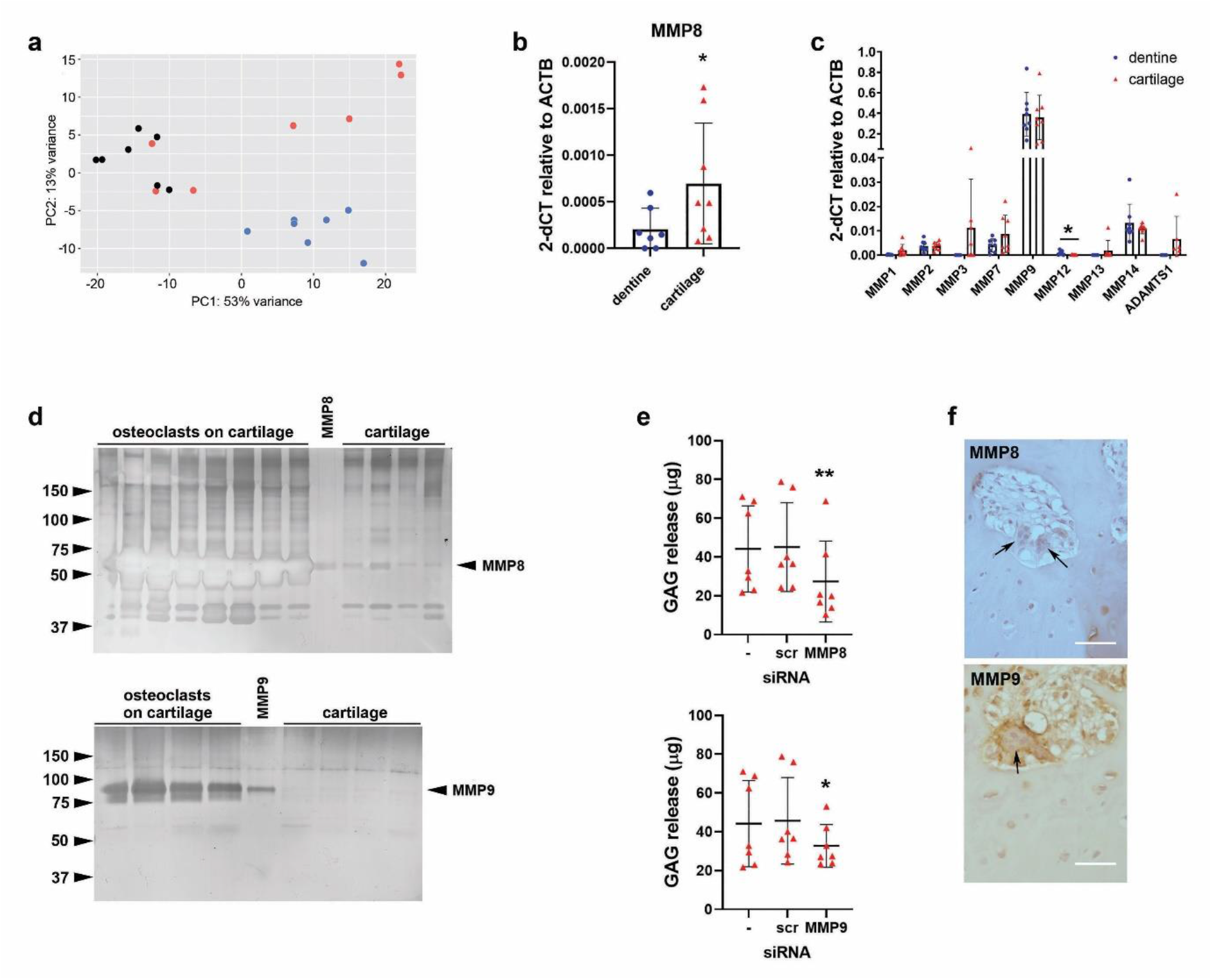
MMP8 and MMP9 drive osteoclast-mediated release of GAG from cartilage. (a) Principal Component Analysis (PCA) plot of osteoclasts on acellular cartilage (red), dentine (blue) and cell culture plastic (black), n=7. (b, c) RT-qPCR analysis of differences in expression of MMPs by osteoclasts differentiated on dentine or acellular cartilage: (b) MMP8, n=8; (c) MMP1, 2, 3, 7, 9, 12, 13, 14 and ADAMTS1. n=8. (d) Representative zymography of MMP8 and MMP9 from the conditioned media of osteoclasts differentiated on acellular cartilage compared to recombinant protein and cartilage alone. (e) GAG released by osteoclasts differentiated on acellular cartilage and transfected with siRNA targeting MMP8 or MMP9 on day 7 of differentiation, versus no siRNA (-) and scrambled siRNA (scr) controls, n=7. (f) Human OA tissue sections stained for MMP8 (top) and MMP9 (bottom). Arrows indicate multinucleated osteoclasts. Scale bar = 100 μm. *, p<0.05; **, p<0.01

**Table 1:**
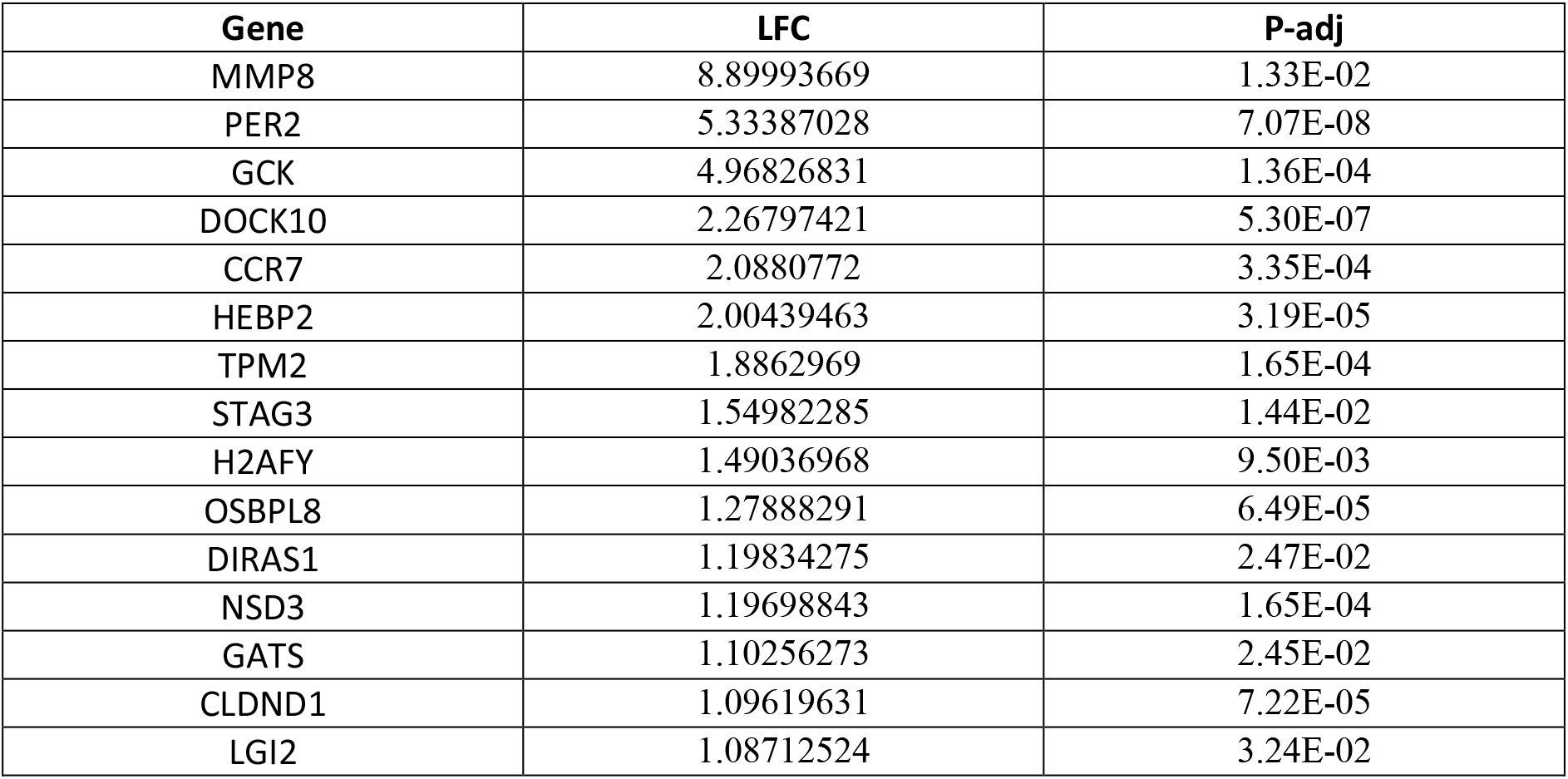
Top 15 over-expressed genes in osteoclasts on cartilage. LFC, Log fold change; padj, p value adjusted.

Gelatin zymography confirmed production of functional MMP8 and MMP9 by osteoclasts cultured on cartilage (Figure 4d). Isoform-specific siRNA-mediated knockdown of MMP8 or MMP9 (Supplementary Figure 4a) reduced osteoclast-mediated GAG release from cartilage explants by 39% and 28% respectively (Figure 4e), without affecting GAG release from cartilage alone (Supplementary Figure 4b). Immunohistochemistry of human OA tissue confirmed strong expression of MMP8 and MMP9 in osteoclasts located between the articular cartilage and subchondral bone (Figure 4f). Chondrocytes surrounded by the cartilage ECM expressed lower levels of both MMPs.

## Discussion

This manuscript identifies MMP8 and MMP9 as the soluble MMPs primarily responsible for osteoclast-mediated degradation of cartilage. It also suggests a role for interactions between osteoclasts and chondrocytes in maintaining joint function. We suggest that, in the healthy joint, chondrocytes and bone-resident osteoclasts exert distant effects on the formation and / or activity of the other cell type in order to maintain joint integrity. However, when the joint architecture is compromised, in conditions such as RA and OA, chondrocytes enhance osteoclast formation within cartilage which drives pathological degradation of the cartilage matrix.

Osteoclasts differentiated on cartilage explants formed large multi-nucleated cells with the equivalent immunophenotypic characteristics (TRAP+, VNR+) and gene expression profile (*CTSK, MMP9, ATP6V1A 21S, CAII, CLCN7, STAT3*) of osteoclasts on dentine. Despite these similarities, osteoclasts on cartilage had a more rounded three-dimensional profile and did not form the F-actin rings characteristic of the sealing zone. Human osteoclasts on cartilage in RA also form a less visible ruffled border without actin rings [28]. It may be that minerals are necessary to induce formation of the sealing zone. Murine osteoclasts cultured on glass coverslips half-coated with apatite only form a sealing zone on the mineralized surface [29]. Without a sealing zone, the ruffled border cannot form, and no resorption pits will be produced. Rabbit osteoclasts form resorption pits on human femoral cortical bone but not on demineralised bone, again suggesting that osteoclasts do not polarise in the absence of minerals [30]. The absence of a ruffled border and sealing zone in osteoclasts on cartilage is probably due to differences in its mechanical, physical and chemical properties compared to dentine and suggests that acidification is not necessary to degrade cartilage, as supported by the lack of effect of bafilomycin on osteoclast-mediated release of GAG.

Despite the absence of resorption pits, clear evidence of cartilage matrix degradation was evident by SEM. Crumbling of cartilage matrix and fragmentation of collagen fibrils adjacent to osteoclasts on cartilage is also evident in the mandibles of rat foetuses [3]. Matrix degradation was confirmed by the increased release of cartilage matrix GAG in the presence of osteoclasts, as we described previously [18]. This was observed despite the high level of natural inter-individual variation in both the resorption / digestion capacity of primary human osteoclasts and the basal rate of cartilage degradation between cartilage donors (Figure 2a).

This is the first study to identify the specific MMPs responsible for cartilage degradation by osteoclasts. Osteoclasts cultured on cartilage secreted active forms of both MMP8 and MMP9 and inhibition of expression of either MMP with siRNA inhibited the degradation of proteoglycans from cartilage by osteoclasts. Only Lovfall et al [21] have previously described how inhibition of pathways important for osteoclast-mediated resorption of bone affects their ability to degrade cartilage. In agreement with our study, the pan-MMP inhibitor GM6001 inhibited osteoclast-mediated degradation of cartilage, whereas inhibitors of osteoclast-mediated bone resorption (cathepsin K inhibitors, vacuolar-type H+-ATPase (V-ATPase) inhibitors) had no effect [21,31]. The lack of effect of V-ATPase inhibitors on cartilage degradation by osteoclasts supports the hypothesis derived from microscopy studies that acidification is not necessary for osteoclasts to degrade cartilage. Support for our data also comes from work on bone; MMP inhibitors only reduce osteoclast-mediated release of collagen from bone when the bone is decalcified [32]. Immunohistochemistry demonstrated expression of MMP8 and MMP9 in osteoclasts in contact with cartilage in OA patient tissue, as we have previously described for MMP9 [18]. Osteoclasts invading the articular cartilage in knee OA also express MMP1, MMP3 and MMP13 [7], suggesting that other MMPs might also contribute to osteoclast-mediated cartilage degradation.

MMP8 was investigated on the basis of its overexpression in osteoclasts cultured on cartilage versus dentine. This is the first time that the gene expression profile of osteoclasts has been compared when they are in contact with their two primary natural substrates. Most osteoclast studies are performed on tissue culture plastic or glass [33] and fail to address the critical role of the substrate in osteoclast differentiation, polarisation and activation. The few publications to study effects of the bone substrate on osteoclastogenesis have performed comparisons with osteoclasts cultured on synthetic cell culture plastic [34,35]. We found no difference in the expression of classical osteoclast genes on a bone versus cartilage substrate, an observation mirrored in murine bone marrow macrophages (BMM) differentiated into osteoclasts on devitalised mouse bone versus plastic [35]. Differential expression of non-classical osteoclast genes could explain the different cytoskeletal arrangements observed on bone versus cartilage. Crotti et al found that expression of annexin A8 increases in murine BMM-derived osteoclasts cultured on bone versus plastic, which regulates cytoskeletal reorganization and formation of the F-actin ring in osteoclasts generated on a mineralised matrix [34]. Principal components analysis revealed osteoclasts on plastic to cluster, to some extent, with osteoclasts on cartilage and away from osteoclasts on bone, suggestive of a broad distinction between the gene expression profile of osteoclasts on their native bone substrate and those in non-bone environments.

The distinction between bone-resident and non-bone-resident osteoclasts becomes evident when considering interactions between chondrocytes and osteoclasts. Bone-resident osteoclasts exerted a protective effect on the cellular, but not acellular, cartilage matrix, suggestive of distant effects of an unidentified osteoclast secreted product(s) on active chondrocytes. Similarly, chondrocytes inhibited the number of distant osteoclasts formed on bone. However, when osteoclasts were cultured on plastic (a ‘non-bone’ substrate) these osteoclasts exerted no indirect, protective effect on cartilage matrix and chondrocytes promoted their formation. This suggests a potential homeostatic relationship between osteoclasts and chondrocytes that maintains joint integrity in the normal joint but drives matrix degradation in disease conditions where joint integrity has been lost.

Some insight into this homeostatic relationship can be obtained from OA. As disease progresses in murine models of OA, increased diffusion of small molecules between bone and cartilage allows direct crosstalk between the cell types due to mechanisms including the penetration of blood vessels and microcracks into the calcified cartilage and increased hydraulic conductance of the articular cartilage and subchondral bone plate [36-38]. Pathways implicated in the general crosstalk between cartilage and bone in OA include the TGF-β/Smad, Wnt/β-catenin, RANK/RANKL/OPG, and MAPK pathways [39]. Considering effects on osteoclasts, the RANK/RANKL/OPG pathway is potentially most interesting. RANKL expression is increased in human OA cartilage compared with normal cartilage, largely due to its expression by hypertrophic OA chondrocytes [40]. In equine OA, the relationship between RANKL expression in the articular cartilage and osteoclast density is stronger than in the subchondral bone and correlates with the number of microcracks, suggesting that cartilage RANKL recruits osteoclasts which contribute to cartilage degradation [41]. In C57BL/6J mice with surgically induced OA, exogenous OPG, a natural decoy receptor for RANKL and osteoclast inhibitor, protected articular cartilage from the progression of OA and also prevented chondrocyte apoptosis [14]. Further evidence that osteoclasts and chondrocytes cooperate to drive OA comes from experiments using osteoclast-targeting bisphosphonates. Alendronate, primarily used to target osteoclast-mediated bone resorption, is chondroprotective in murine models of OA, inhibiting vascular invasion of calcified cartilage and increasing cartilage thickness [12,42].

In summary, our data demonstrates that osteoclasts can perform proteolysis-driven cartilage degradation and that MMP8 and MMP9 are involved in this process. It also provides new insights into the complexity of the interactions between bone and cartilage; specifically, regarding how interactions between osteoclasts and chondrocytes depend on whether osteoclasts reside on bone or a non-bone substrate. This data is relevant to a variety of pathological joint conditions where understanding and controlling the activity of osteoclasts presents an attractive option for targeting cartilage degradation.

## Acknowledgements

HJK was supported by Arthritis Research UK (MP/19200) and the Rosetrees Trust (A1726). SS was funded by the National Institute of Health Research (NIHR) Oxford Biomedical Research Centre. APC was funded by a Medical Research Council (MRC) Career Development Award. Work in the Nuffield Department of Orthopaedics, Rheumatology and Musculoskeletal Sciences (NDORMS) is additionally supported by the (NIHR) Oxford Biomedical Research Centre. SEM pictures were taken with the guidance of Dr Kalin Dragnevski, Department of Engineering Science, University of Oxford, Oxford, UK. We would also like to thank Louise Hill, Dr Fiona Watt and the Oxford Musculoskeletal Biobank for the collection of human cartilage.

## References

1 Staines, K. A., Pollard, A. S., McGonnell, I. M., Farquharson, C. & Pitsillides, A. A. Cartilage to bone transitions in health and disease. J Endocrinol 219, R1–R12, doi:10.1530/JOE-13-0276 (2013).

2 Schenk, R. K., Spiro, D. & Wiener, J. Cartilage resorption in the tibial epiphyseal plate of growing rats. J Cell Biol 34, 275–291, doi:10.1083/jcb.34.1.275 (1967).

3 Savostin-Asling, I. & Asling, C. W. Transmission and scanning electron microscope studies of calcified cartilage resorption. Anat Rec 183, 373–391, doi:10.1002/ar.1091830303 (1975).

4 Ota, N. et al. Accelerated cartilage resorption by chondroclasts during bone fracture healing in osteoprotegerin-deficient mice. Endocrinology 150, 4823–4834, doi:10.1210/en.2009-0452 (2009).

5 Bromley, M. & Woolley, D. E. Chondroclasts and osteoclasts at subchondral sites of erosion in the rheumatoid joint. Arthritis Rheum 27, 968–975 (1984).

6 Bromley, M., Bertfield, H., Evanson, J. M. & Woolley, D. E. Bidirectional erosion of cartilage in the rheumatoid knee joint. Ann Rheum Dis 44, 676–681 (1985).

7 Shibakawa, A. et al. The role of subchondral bone resorption pits in osteoarthritis: MMP production by cells derived from bone marrow. Osteoarthritis Cartilage 13, 679–687, doi:10.1016/j.joca.2005.04.010 (2005).

8 Luo, G., Li, F., Li, X., Wang, Z. G. & Zhang, B. TNFalpha and RANKL promote osteoclastogenesis by upregulating RANK via the NFkappaB pathway. Mol Med Rep 17, 6605–6611, doi:10.3892/mmr.2018.8698 (2018).

9 Lin, Y. J., Anzaghe, M. & Schulke, S. Update on the Pathomechanism, Diagnosis, and Treatment Options for Rheumatoid Arthritis. Cells 9, doi:10.3390/cells9040880 (2020).

10 Herrak, P. et al. Zoledronic acid protects against local and systemic bone loss in tumor necrosis factor-mediated arthritis. Arthritis Rheum 50, 2327–2337, doi:10.1002/art.20384 (2004).

11 Cohen, S. B. et al. Denosumab treatment effects on structural damage, bone mineral density, and bone turnover in rheumatoid arthritis: a twelve-month, multicenter, randomized, double-blind, placebo-controlled, phase II clinical trial. Arthritis Rheum 58, 1299–1309 (2008).

12 Hayami, T. et al. The role of subchondral bone remodeling in osteoarthritis: reduction of cartilage degeneration and prevention of osteophyte formation by alendronate in the rat anterior cruciate ligament transection model. Arthritis Rheum 50, 1193–1206, doi:10.1002/art.20124 (2004).

13 Kadri, A. et al. Inhibition of bone resorption blunts osteoarthritis in mice with high bone remodelling. Ann Rheum Dis 69, 1533–1538, doi:10.1136/ard.2009.124586 (2010).

14 Shimizu, S. et al. Prevention of cartilage destruction with intraarticular osteoclastogenesis inhibitory factor/osteoprotegerin in a murine model of osteoarthritis. Arthritis Rheum 56, 3358–3365, doi:10.1002/art.22941 (2007).

15 Blangy, A. et al. The osteoclast cytoskeleton - current understanding and therapeutic perspectives for osteoporosis. J Cell Sci 133, doi:10.1242/jcs.244798 (2020).

16 Odgren, P. R., Witwicka, H. & Reyes-Gutierrez, P. The cast of clasts: catabolism and vascular invasion during bone growth, repair, and disease by osteoclasts, chondroclasts, and septoclasts. Connect Tissue Res 57, 161–174, doi:10.3109/03008207.2016.1140752 (2016).

17 Gravallese, E. M. et al. Identification of cell types responsible for bone resorption in rheumatoid arthritis and juvenile rheumatoid arthritis. Am J Pathol 152, 943–951 (1998).

18 Knowles, H. J. et al. Chondroclasts are mature osteoclasts which are capable of cartilage matrix resorption. Virchows Arch 461, 205–210, doi:10.1007/s00428-012-1274-3 (2012).

19 Komuro, H. et al. The osteoprotegerin/receptor activator of nuclear factor kappaB/receptor activator of nuclear factor kappaB ligand system in cartilage. Arthritis Rheum 44, 12, 2768–2776, doi:10.1002/1529-0131(200112) (2001).

20 Nakao, K. et al. Collaborative action of M-CSF and CTGF/CCN2 in articular chondrocytes: possible regenerative roles in articular cartilage metabolism. Bone 36, 884–892, doi:10.1016/j.bone.2004.10.015 (2005).

21 Lofvall, H. et al. Osteoclasts degrade bone and cartilage knee joint compartments through different resorption processes. Arthritis Res Ther 20, 67, doi:10.1186/s13075-018-1564-5 (2018).

22 Mahoney, D. J. et al. TSG-6 inhibits osteoclast activity via an autocrine mechanism and is functionally synergistic with osteoprotegerin. Arthritis Rheum 63, 1034–1043, doi:10.1002/art.30201 (2011).

23 Oshita, K. et al. Human mesenchymal stem cells inhibit osteoclastogenesis through osteoprotegerin production. Arthritis Rheum 63, 1658–1667, doi:10.1002/art.30309 (2011).

24 Cribbs APL.-V.S., George C et al. CGAT-core: a python framework for building scalable, reproducible computational biology workflows. [version 2; peer review: 1 approved, 1 approved with reservations]. F1000Research 2019, 8:377 (https://doi.org/10.12688/f1000research.18674.2)

25 Sims, D. et al. CGAT: computational genomics analysis toolkit. Bioinformatics 30, 1290–1291, doi:10.1093/bioinformatics/btt756 (2014).

26 Dobin, A. et al. STAR: ultrafast universal RNA-seq aligner. Bioinformatics 29, 15–21, doi:10.1093/bioinformatics/bts635 (2013).

27 Love, M. I., Huber, W. & Anders, S. Moderated estimation of fold change and dispersion for RNA-seq data with DESeq2. Genome Biol 15, 550, doi:10.1186/s13059-014-0550-8 (2014).

28 Bromley, M. & Woolley, D. E. Histopathology of the rheumatoid lesion. Identification of cell types at sites of cartilage erosion. Arthritis Rheum 27, 857–863 (1984).

29 Saltel, F., Destaing, O., Bard, F., Eichert, D. & Jurdic, P. Apatite-mediated actin dynamics in resorbing osteoclasts. Mol Biol Cell 15, 5231–5241, doi:10.1091/mbc.e04-06-0522 (2004).

30 Chambers, T. J., Thomson, B. M. & Fuller, K. Effect of substrate composition on bone resorption by rabbit osteoclasts. J Cell Sci 70, 61–71 (1984).

31 Neutzsky-Wulff, A. V. et al. Alterations in osteoclast function and phenotype induced by different inhibitors of bone resorption--implications for osteoclast quality. BMC Musculoskelet Disord 11, 109, doi:10.1186/1471-2474-11-109 (2010).

32 Henriksen, K. et al. Degradation of the organic phase of bone by osteoclasts: a secondary role for lysosomal acidification. J Bone Miner Res 21, 58–66, doi:10.1359/JBMR.050905 (2006).

33 Sharma, S. M. et al. Genetics and genomics of osteoclast differentiation: integrating cell signaling pathways and gene networks. Crit Rev Eukaryot Gene Expr 16, 253–277, doi:10.1615/critreveukargeneexpr.v16.i3.40 (2006).

34 Crotti, T. N. et al. Bone matrix regulates osteoclast differentiation and annexin A8 gene expression. J Cell Physiol 226, 3413–3421, doi:10.1002/jcp.22699 (2011).

35 Purdue, P. E. et al. Comprehensive profiling analysis of actively resorbing osteoclasts identifies critical signaling pathways regulated by bone substrate. Sci Rep 4, 7595, doi:10.1038/srep07595 (2014).

36 Hwang, J. et al. Increased hydraulic conductance of human articular cartilage and subchondral bone plate with progression of osteoarthritis. Arthritis Rheum 58, 3831–3842, doi:10.1002/art.24069 (2008).

37 Pan, J. et al. Elevated cross-talk between subchondral bone and cartilage in osteoarthritic joints. Bone 51, 212–217, doi:10.1016/j.bone.2011.11.030 (2012).

38 Botter, S. M. et al. Osteoarthritis induction leads to early and temporal subchondral plate porosity in the tibial plateau of mice: an in vivo microfocal computed tomography study. Arthritis Rheum 63, 2690–2699, doi:10.1002/art.30307 (2011).

39 Zhou, X., Cao, H., Yuan, Y. & Wu, W. Biochemical Signals Mediate the Crosstalk between Cartilage and Bone in Osteoarthritis. Biomed Res Int 2020, 5720360, doi:10.1155/2020/5720360 (2020).

40 Kwan Tat, S. et al. Modulation of OPG, RANK and RANKL by human chondrocytes and their implication during osteoarthritis. Rheumatology (Oxford) 48, 1482–1490, doi:10.1093/rheumatology/kep300 (2009).

41 Bertuglia, A. et al. Osteoclasts are recruited to the subchondral bone in naturally occurring post-traumatic equine carpal osteoarthritis and may contribute to cartilage degradation. Osteoarthritis Cartilage 24, 555–566, doi:10.1016/j.joca.2015.10.008 (2016).

42 Bikle, D. D. et al. Alendronate increases skeletal mass of growing rats during unloading by inhibiting resorption of calcified cartilage. J Bone Miner Res 9, 1777–1787, doi:10.1002/jbmr.5650091115 (1994).

