## Supplementary for "Loss of mutually protective effects between osteoclasts and chondrocytes in damaged joints drives osteoclast-mediated cartilage degradation via matrix metalloproteinases"

**Supplementary table 1**

| Gene name | Forward primer | Reverse primer |
| --- | --- | --- |
| ACP5 | TGAGGACGTATTCTCTGACCG | CACATTGGTCTGTGGGATCTTG |
| ATP6V1A 21S | CCATCGGAACTACCATGCAGG | TCCACAGAAGAGGTTAGACAGG |
| CA2 | GGACAAAGCAGCTTCACTTGG | CCTAGAACGGCCAGTCCATC |
| CLCN7 | CTAAGAAGGTGTCCTGGTCCG | GGAGGGTCCATATCCTGGCG |
| CTSK | GCAGAAGAACCGGGGTATTGA | GAAGGAGGTCAGGCTTGCAT |
| MMP2 | TACAGGATCATTGGCTACACACC | GGTCACATCGCTCCAGACT |
| MMP7 | GAGTGAGCTACAGTGGGAACA | CTATGACGCGGGAGTTTAACAT |
| MMP12 | AACCAACGCTTGCCAAATCC | TTTCCCACGGTAGTGACAGC |
| MMP14 | ATCTGCCTCTGCCTCACCTA | AAGCCCCATCCAAGGCTAAC |

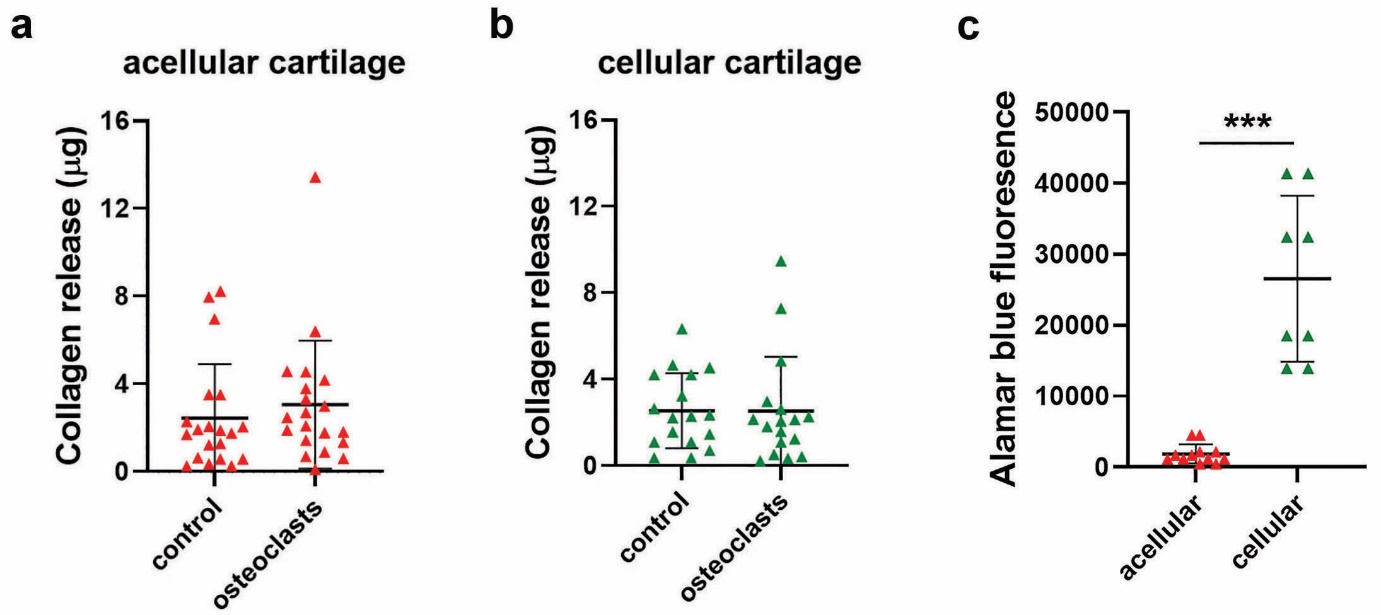

**Supplementary Figure 1:** **Substrate-dependent regulation of osteoclasts by chondrocytes.** (a, b) Osteoclast-mediated release of collagen from (a) acellular (n=20) or (b) cellular (n=17) cartilage after 13 days of differentiation. (c) Alamar blue fluorescence of acellular and cellular cartilage on day 13 of culture (no osteoclasts, n=8). ***, p<0.0001.

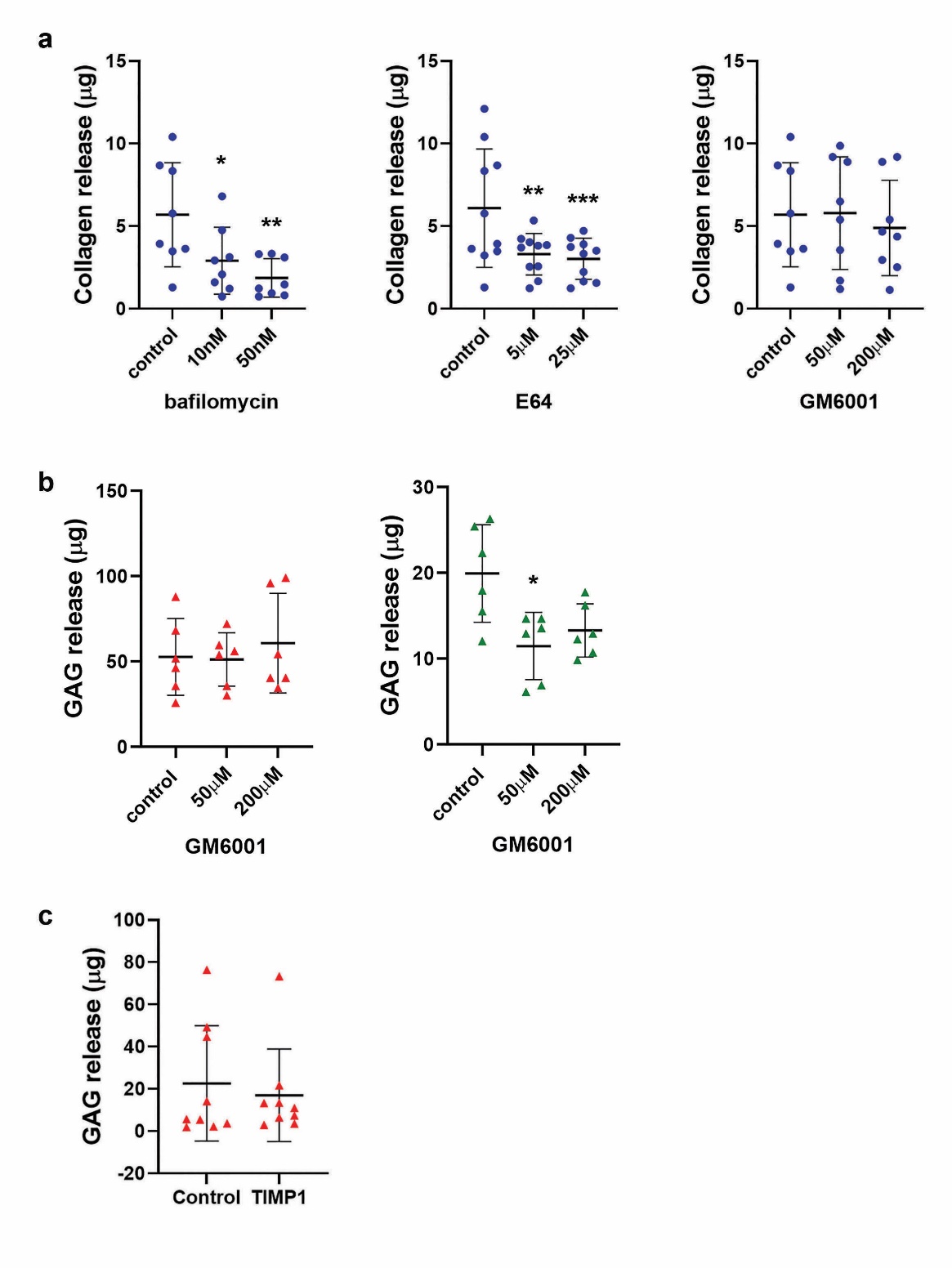

**Supplementary Figure 2:** **MMPs drive osteoclast-mediated degradation of cartilage.** (a) Effect of bafilomycin (n=8), E64 (n=10) and GM6001 (n=8) on osteoclast-mediated release of collagen from dentine after 13 days of differentiation. (b) Effect of GM6001 on basal release of GAG from acellular (red) and cellular (green) cartilage in the absence of osteoclasts, n=6. (c) Effect of recombinant human TIMP1 (100 nm) on basal release of GAG from acellular cartilage in the absence of osteoclasts, n=9. *, p<0.05; ***, p<0.01; ***, p<0.0001.

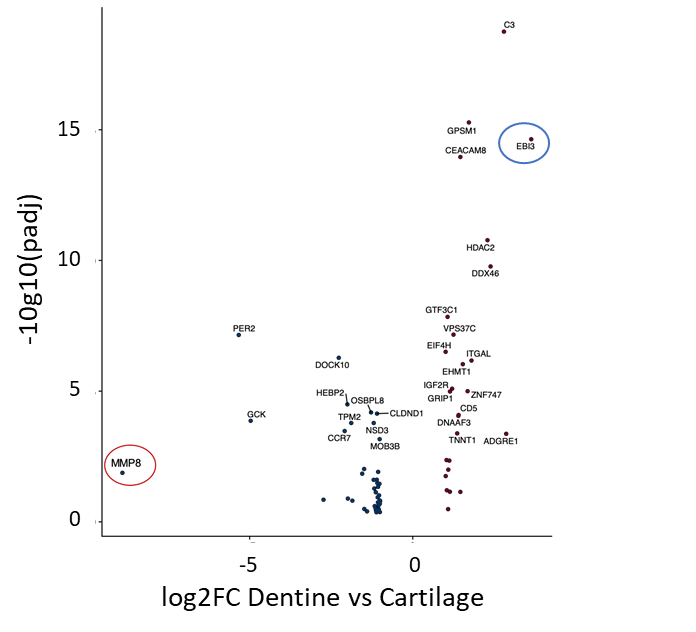

**Supplementary Figure 3: Volcano plot showing genes overexpressed in osteoclasts cultured on dentine versus cartilage.** Genes overexpressed on cartilage are on the left of the plot, those overexpressed on dentine are on the right. FC, fold change; padj, p value adjusted.

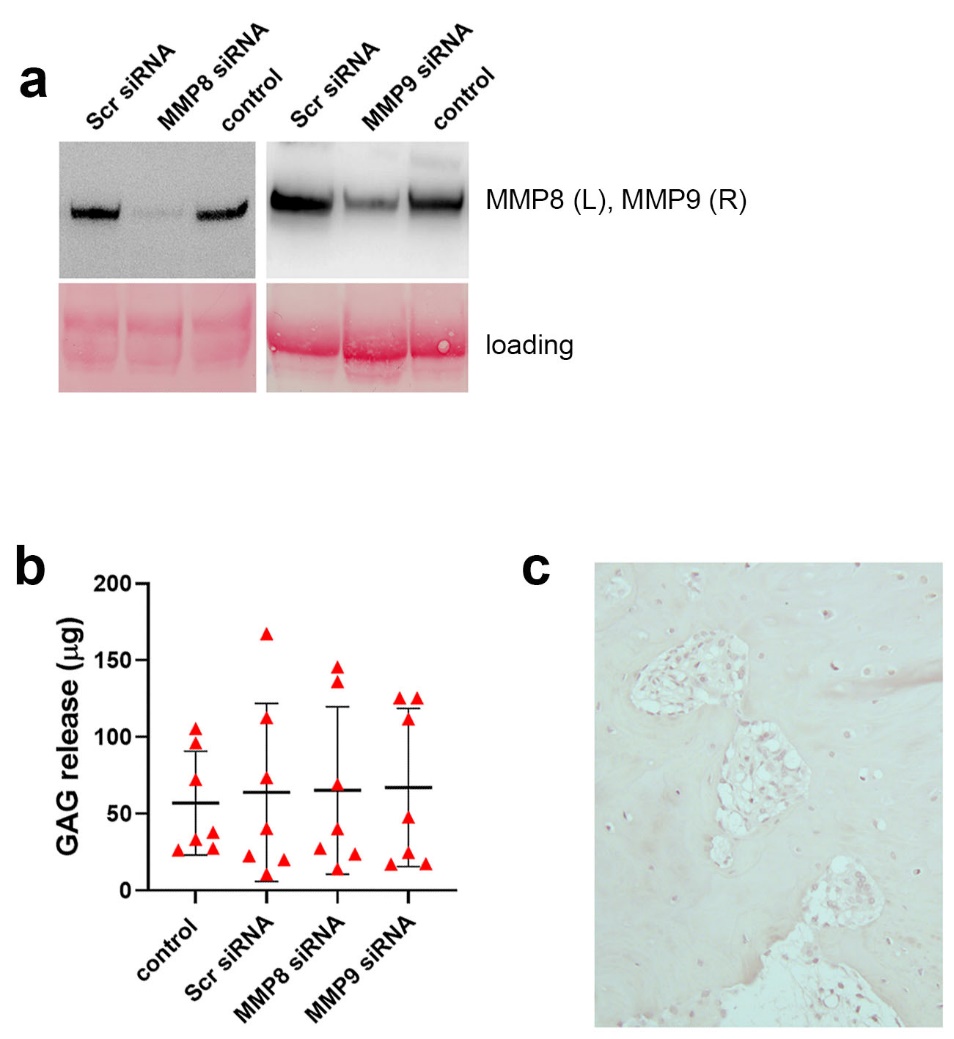

**Supplementary Figure 4:** **MMP8 and MMP9 are the primary MMPs responsible for osteoclast-mediated release of GAG from cartilage.** (a) Western blots showing MMP8 and MMP9 expression in

osteoclasts following treatment with MMP8- or MMP9-siRNA or a scrambled siRNA control. Loading control = Ponceau staining. (b) Effect of MMP8 or MMP9 siRNA on basal release of GAG from acellular cartilage in the absence of osteoclasts, n=7. (c) IgG negative control immunohistochemistry of OA tissue section.
